# In search of visual opsin of *Lissachatina fulica*: *in silico* prediction

**DOI:** 10.1101/2024.03.19.585790

**Authors:** Irina N. Dominova, Daniil A. Fedotov, Svetlana Shirina, Vladimir Zikin, Valerii V. Zhukov

## Abstract

Here we present a complex study of possible visual opsin of *L. fulica*. 8 candidate sequences were chosen for visual opsin protein and only one (Afu005002) was confirmed as a possible candidate after domain search, structural (structures were stimulated with AlphaFold2) and BLAST analysis. In order to evaluate spectroscopic properties of the Afu005002 protein, we have computed excitation energy and intensity of the main absorption band using QM/MM methodology at the ADC(2) level of theory. The theoretically predicted position of the main absorption maxima shows a good agreement with available experimental data.

## Introduction

Opsins represent a large group of transmembrane proteins, where visual opsins form the molecular basis of light-sensitive pigments [1, 2]. Structurally and functionally, visual opsins are members of the serpentine family of G-protein-coupled proteins (GPCRs). These proteins form 7 transmembrane domains and convert the energy of an absorbed light into a molecular signal for activation of intracellular signalling pathways [3].

Being bound to the chromophore group through Schiff base, visual opsins represent a light-sensitive chromoprotein. The light sensitivity of the visual pigment ranges from ultraviolet to infrared usually with two distinct peaks: the first and the strongest peak and a low intensity non-functional peak shifted to the blue area [4]. The position of the main absorption band reflects the illuminance characteristic of the animalșs habitat. There are only 4 chromophores in the animal kingdom: retinal, 3-dehydroretinal, and 3-and 4-hydroxyretinal [5]. At the same time, there are many visual pigments that differ in the position of the absorption maximum. Therefore, the position of the absorption band of each visual pigment is determined by the amino acid sequence of opsin [6,7] and specifically by its counterions [8,9].

Nowadays, despite significant progress in determination of structure and photochemistry of visual opsins in some animal groups, the overall picture of evolutionary changes remains incomplete. A significant gap is the lack of such information for gastropod mollusks, the largest soft-bodied type in terms of the number of species. *Lissachatina fulica* [10] snail is an example of a promising model organism for its capability of specific body parts regeneration, such as the eye tentacle. The regeneration ability of *Lissachatina fulica* was first demonstrated by Bobkova and co-authors [11].

The investigation of the regeneration processes in the eye tentacle of *Lisachatina fulica* can be beneficial for treatment of such socially significant diseases such as Alzheimer’s and Parkinson’s diseases, consequences of brain injuries, neuroinflammatory processes in the human brain, etc. However, there is no available data on visual opsin of *L. fulica* as well as its structure, pH-dependent spectroscopic characteristics, etc.

In this study, we have performed a theoretical prediction of *Lisachatina fulica* visual opsin through (1) confirmation of the presence of domains characteristic of opsin proteins; (2) determination of the degree of identity of the found sequences with proteins of gastropods, cephalopods and bivalves; (3) construction of 3D structures of the predicted proteins and comparison with the known structure of squid opsin (4) simulation of (pH-dependent) spectroscopic characteristics and its comparison with theoretical data.

## Materials and methods

### Compiling data from databases

For the Opsin protein prediction in *Lissachatina fulica*, we selected only *Gastropoda* classe. Opsin proteins sequences were taken from the NCBI Protein database (https://www.ncbi.nlm.nih.gov/protein/ (accessed on 15 March 2024)) and are presented in Table S1. We excluded proteins with PREDICTED and LOW QUALITY status, as well as hypothetical protein, too short and long sequences from further analysis. Thus, 79 sequences of Opsins (Table S1) were collected for further analysis.

The primary predicted sequences of *L. fulica* proteins were taken from Guo’s article [12] supplementary materials deposited in the GigaDB database (http://gigadb.org/dataset/100647 (accessed on 31 October 2022)).

### Protein prediction

Amino acid sequence of opsins in the *L. fulica* was performed using the HMMER software (version 3.3.2) [HMMER: biosequence analysis using profile hidden Markov models]. To achieve this objective, we performed a multiple sequence alignment of 79 previously gathered rhodopsin molluscan proteins using Unipro UGENE software (version 45.1) developed by Unipro in Russia [13]. The alignment utilized Clustal Omega [14] and MUSCLE [15] algorithms integrated within UGENE. Initially, Clustal Omega was employed for alignment with the following parameters: 100 iterations, maximum number of guide-tree iterations set at 100, and maximum number of Hidden Markov Models (HMM) iterations at 100. Subsequently, the alignment was further refined using MUSCLE with a maximum of 100 iterations, encompassing the entire amino acid sequences.

Utilizing the alignment obtained, we generated an HMM3-profile within Unipro UGENE. Subsequently, this profile was employed to forecast the protein sequences under investigation in *L. fulica* using the hmmsearch algorithm for protein sequence analysis, seamlessly integrated into HMMER. In our study, the target sequence database comprised the primary predicted *L. fulica* protein sequences archived in the GigaDB database. Following prediction, sequences with an E-value below the predefined threshold of 10^-60^ were singled out for further analysis and validation.

### Domain search

The presence of the opsin core domains in the identified *L. fulica* sequences was verified using the InterProScan website (https://www.ebi.ac.uk/interpro/ (accessed on 16 March 2024)) [16].

We also performed multiple sequence alignments using UGENE software, utilizing the Clustal Omega and MUSCLE algorithms with identical parameters to those employed in constructing the HMM-profile. These alignments were aimed at pinpointing and characterizing conserved domains within the predicted sequences. The reference sequences for the conserved domains included pfam00001 “7tm_1” for the 7 transmembrane receptor (rhodopsin family) (https://www.ncbi.nlm.nih.gov/Structure/cdd/cddsrv.cgi?uid=278431/ (accessed on 16 March 2024)) and cd00637 “7tm_classA_rhodopsin-like” for the rhodopsin receptor-like class A family of the seven-transmembrane G protein-coupled receptor superfamily (https://www.ncbi.nlm.nih.gov/Structure/cdd/cddsrv.cgi?uid=410626/ (accessed on 16 March 2024)). The alignment results were visualized using the Jalview software [17].

### BLAST analysis

The predicted proteins of *L. fulica* were scanned using the BLASTp algorithm against gastropod, cephalopod, and bivalve opsin proteins stored in the NCBI GenBank’s protein database of Reference proteins (refseq_protein) (https://blast.ncbi.nlm.nih.gov/Blast.cgi/ (accessed on 17 March 2024)).

### 3D structure

The 3D structure of the predicted proteins was derived by AlphaFold2 Colab, employing MMseqs2 (v1.5.3) (https://colab.research.google.com/github/sokrypton/ColabFold/blob/main/AlphaFold2.ipynb (accessed on 16 March, 2024)) [18]. All parameters were used by default except for the PDB100^*^ template, and the number of cycles was also increased to the maximum, while limiting the recycle_early_stop_tolerance parameter to 0.05 to optimize the usage time. PyMOL version 2.5.7 was utilized for structuring alignment, visualizing the outcomes, and generating figures [19].

### Structure alignment

To select the most likely candidate visual opsin and/or opsins among the predicted Lissachatina fulica proteins, the 3D structures of the predicted Lissachatina fulica proteins obtained using AlphaFold2 were aligned and overlaid with the 3D structure of the squid *Todarodes pacificus* rhodopsin resolved by X-ray structural analysis to 3.70 Å accuracy, taken from the RCSB Protein Data Bank database (https://www.rcsb.org/structure/2ZIY (accessed 25.09.2023)) [20].

3D structure alignment serves not only to ascertain whether the analyzed proteins share similar functions but also facilitates comparisons among proteins with low sequence similarity that fold into analogous forms. This is particularly valuable when evolutionary relationships are challenging to discern through one-dimensional sequence alignment alone.

Structures were compared and aligned using PyMOL software, which adopts a two-step process for structure alignment: initially, sequence alignment is conducted, followed by minimizing the Root Mean Square Deviation (RMSD) between aligned residues. RMSD quantifies the extent to which a given molecular structure diverges from a reference geometry. In bioinformatics, RMSD of atomic positions provides a measure of the average distance between overlapping protein atoms [21].

### AlphaFold2 structure processing

The 3D model predicted with AlphaFold2 was used as an initial structure for a computational model. The protonation states of titratable residues were determined with on-line service propKa [22] Protons were added using the pdb4amber program from AmberTools [23] package. Since AlphaFold2 can not predict the structure of the retinal chromophore, the 11-cis retinal molecule was added manually into Alphafold2 predicted structure. The reason for this choice was due to the high similarity (see Figure 1) of predicted structure and structure of squid rhodopsin (PDB: 2Z73).

**Figure 1.**
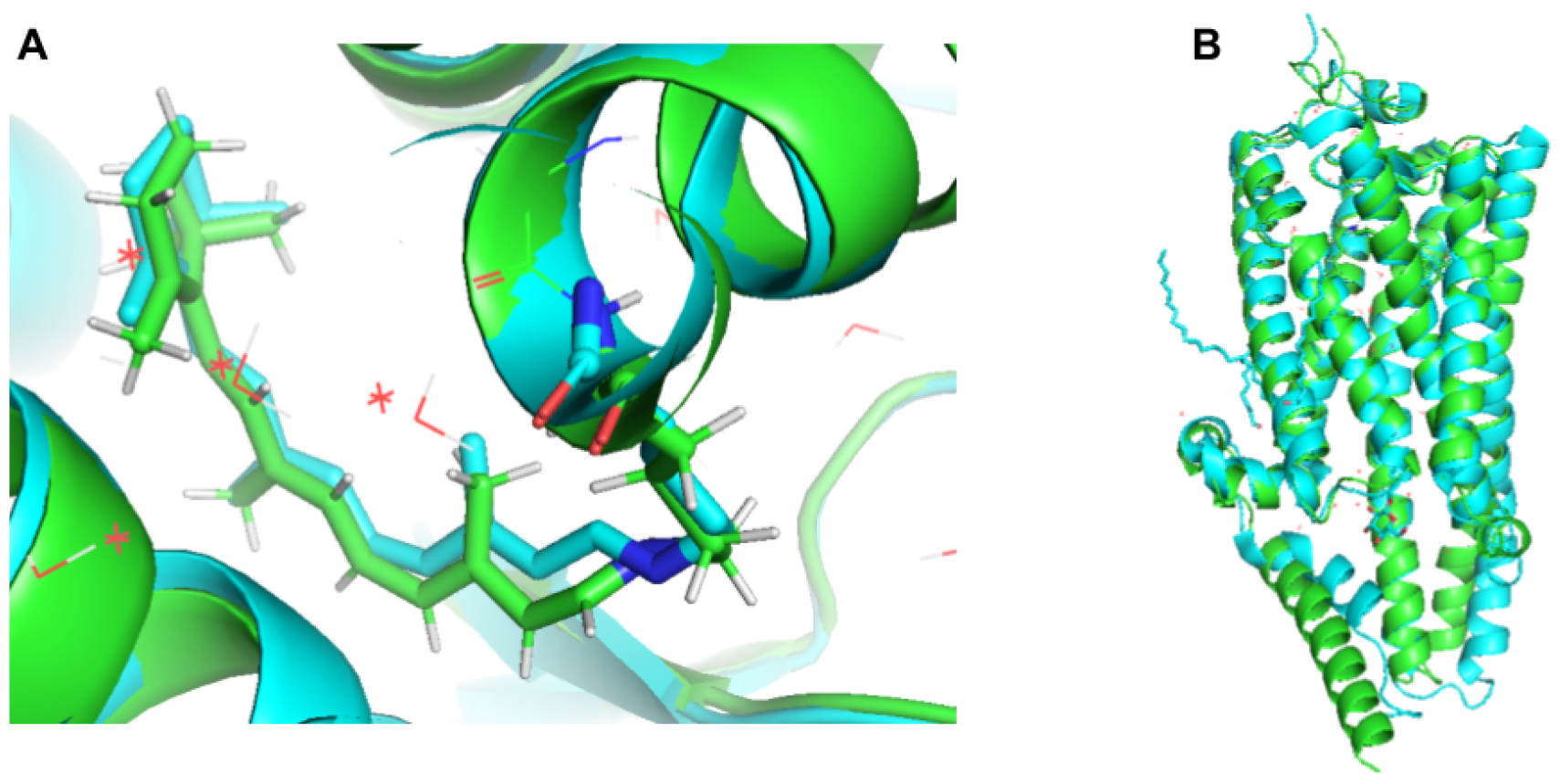
(A) structural comparison of AlphaFold2 predicted protein structure shown in green and structure of squid rhodopsin (PDB: 2Z73) shown in cyan; (B) alignment of both structures (RSMD=0.8).

Then ff14SB AMBER [24] force field parameters were generated with use of the TIP3P water model. [25]. In order to relax the resulting protein structure and resolve possible atom clashes initial energy minimisation was performed at the molecular mechanics (MM) level with ff14SB AMBER force field using Amber package [26.].

### Structure optimization and calculation of excitation energies

Structural optimisation was performed with use of Quantum Mechanics/Molecular Mechanics hybrid [27] method (QM/MM) employing L-BFGS [28] algorithm. The QM part was determined as a retinal protonated Schiff base (PSB) with the part of lysine side chain and, if specified, a sidechain of glutamic residue 175 (GLU175). The QM part was treated with TD-DFT [29,30] (B3LYP) level of theory, appalling Grimme [31] dispersion correction with Becke-Johnson [32 damping, adopting the correlation-consistent cc-pVDZ [33] atomic basis set. For both Hartree-Fock and TD-DFT calculations, the Resolution-of-Identity [34] approximation was used, the auxiliary basis set was chosen equal to the atomic one. The MM part comprised all atoms of the amino acid side chains and water molecules within 5 Å of retinal chromophore. The MM part was treated with ff14SB AMBER force field, while the rest of the protein was frozen. For all protonation models the one-photon absorption spectrum was computed using second order Algebraic Diagrammatic Construction (ADC(2)) method with Dunning’s correlation-consistent basis set ??-pVDZ in Turbomole [36] v7.3 .

## Results and discussion

### Prediction and validation of opsin proteins

The application of HMMER for predicting opsin proteins revealed the existence of 8 sequences of such nature within *L. fulica* (Table 1).

**Table 1.** Significant results (E-value < 10^-60^) of the HMMER analysis for the prediction of Opsin proteins in *L. fulica*.

| Sequence ID | Full sequence | Best 1 domain |
| --- | --- | --- |

|  | E-value | Score | E-value | Score |
| --- | --- | --- | --- | --- |
| Afu005002 | $6.7 \times 10^{-137}$ | 457.6 | $8.2 \times 10^{-137}$ | 457.3 |
| Afu017437 | $2.4 \times 10^{-111}$ | 373.3 | $5.2 \times 10^{-66}$ | 223.8 |
| Afu017435 | $2.8 \times 10^{-105}$ | 353.3 | $3.5 \times 10^{-59}$ | 201.3 |
| Afu004575 | $7.1 \times 10^{-90}$ | 302.5 | $9.2 \times 10^{-72}$ | 242.8 |
| Afu000836 | $6.5 \times 10^{-72}$ | 243.3 | $8.4 \times 10^{-72}$ | 242.9 |
| Afu017436 | $5.7 \times 10^{-64}$ | 217.1 | $6.1 \times 10^{-64}$ | 217.0 |
| Afu004794 | $1.6 \times 10^{-63}$ | 215.7 | $3.2 \times 10^{-63}$ | 214.6 |
| Afu004211 | $1.1 \times 10^{-62}$ | 212.8 | $1.4 \times 10^{-62}$ | 212.6 |

To confirm if the 8 predicted sequences corresponded to opsin proteins, a search for the most common opsin domain was conducted using the InterPro web service (Table 2).

**Table 2.** Results of the domain search in the predicted sequences of *L. fulica* using the InterPro web service.

| InterPro entries<br>- Family | Protein<br>Database | Protein<br>Database<br>ID | Domain | Domain<br>position,<br>amino acid<br>residues | E-value |
| --- | --- | --- | --- | --- | --- |
| Afu005002 |  |  |  |  |  |
| IPR000276 - G<br>protein-coupled<br>receptor,<br>rhodopsin-like | SMART | SM01381 | 7TM_GPCR_Srsx_2 | 59 - 343 | $2.3 \times 10^{-4}$ |
| | Pfam | PF00001 | 7 transmembrane receptor<br>(rhodopsin family) | 65 - 328 | $3.5 \times 10^{-55}$ |
| | PRINTS | PR00237 | Rhodopsin-like GPCR<br>superfamily signature | 272 - 296<br>310 - 336<br>129 - 151<br>165 - 186<br>83 - 104<br>50 - 74<br>215 - 238 | $4.7 \times 10^{-33}$ |
| IPR001760 -<br>Opsin | PRINTS | PR00238 | Opsin signature | 187 - 199<br>307 - 319<br>71 - 83 | $2.7 \times 10^{-8}$ |
| IPR001391 -<br>Opsin lateral<br>eye type | PRINTS | PR00578 | Lateral eye opsin signature | 37 - 51<br>98 - 110<br>337 - 348<br>357 - 370 | $7.0 \times 10^{-5}$ |

| Afu017437 |  |  |  |  |  |
| --- | --- | --- | --- | --- | --- |
| IPR000276 - G protein-coupled receptor, rhodopsin-like | Pfam | PF00001 | 7 transmembrane receptor (rhodopsin family) | 750 - 976 | $4.3 \times 10^{-11}$ |
|  | ProSitePatterns | PS00237 | G-protein coupled receptors family 1 signature. | 250 - 266<br>813 - 829 | - |
| | Pfam | PF00001 | 7 transmembrane receptor (rhodopsin family) | 185 - 438 | $9.3 \times 10^{-12}$ |
| IPR017452 - GPCR, rhodopsin-like, 7TM | ProSiteProfiles | PS50262 | G-protein coupled receptors family 1 profile. | 185 - 458<br>748 - 965 | - |
| - | CDD | cd14978 | 7tmA_FMRFamide_R-like | 173 - 469 | $3.6 \times 10^{-29}$ |
| Afu017435 |  |  |  |  |  |
| IPR000276 - G protein-coupled receptor, rhodopsin-like | PRINTS | PR00237 | Rhodopsin-like GPCR superfamily signature | 921 - 945<br>138 - 162<br>968 - 994<br>212 - 234<br>170 - 191 | $2.5 \times 10^{-6}$ |
|  | ProSitePatterns | PS00237 | G-protein coupled receptors family 1 signature. | 218 - 234<br>782 - 798 | - |
| | Pfam | PF00001 | 7 transmembrane receptor (rhodopsin family) | 153 - 373 | $3.9 \times 10^{-9}$ |
| | | | | 719 - 980 | $7.1 \times 10^{-9}$ |
| IPR017452 - GPCR, rhodopsin-like, 7TM | ProSiteProfiles | PS50262 | G-protein coupled receptors family 1 profile. | 717 - 986<br>153 - 370 | - |
| Afu004575 |  |  |  |  |  |
| IPR000276 - G protein-coupled receptor, rhodopsin-like | PRINTS | PR00237 | Rhodopsin-like GPCR superfamily signature | 107 - 128<br>74 - 98<br>188 - 209<br>522 - 546<br>238 - 261<br>152 - 174<br>560 - 586 | $1.6 \times 10^{-30}$ |
| | Pfam | PF00001 | 7 transmembrane receptor (rhodopsin family) | 89 - 273 | $3.9 \times 10^{-35}$ |
|  | ProSitePro files | PS00237 | G-protein coupled receptors family 1 signature. | 158 - 174 | - |
| IPR017452 - GPCR, rhodopsin-like, 7TM | ProSitePro files | PS50262 | G-protein coupled receptors family 1 profile. | 89 - 578 | - |
| - | PANTHER | PTHR24240 | OPSIN | 61 - 595 | $1.5 \times 10^{-67}$ |
| Afu000836 |  |  |  |  |  |
| IPR000276 - G protein-coupled receptor, rhodopsin-like | PRINTS | PR00237 | Rhodopsin-like GPCR superfamily signature | 28 - 52<br>245 - 269<br>184 - 207<br>286 - 312<br>102 - 124 | $8.9 \times 10^{-6}$ |
| | Pfam | PF00001 | 7 transmembrane receptor (rhodopsin family) | 44 - 277 | $2.9 \times 10^{-14}$ |
|  | ProSitePro files | PS00237 | G-protein coupled receptors family 1 signature. | 108 - 124 | - |
| IPR017452 - GPCR, rhodopsin-like, 7TM | ProSitePro files | PS50262 | G-protein coupled receptors family 1 profile. | 43 - 304 | - |
| Afu017436 |  |  |  |  |  |
| IPR000276 - G protein-coupled receptor, rhodopsin-like | ProSitePatterns | PS00237 | G-protein coupled receptors family 1 signature. | 113 - 129 | - |
| | Pfam | PF00001 | 7 transmembrane receptor (rhodopsin family) | 48 - 304 | $2.7 \times 10^{-11}$ |
| | PRINTS | PR00237 | Rhodopsin-like GPCR superfamily signature | 256 - 280<br>33 - 57<br>107 - 129<br>303 - 329 | $7.8 \times 10^{-6}$ |
| IPR017452 - GPCR, rhodopsin-like, 7TM | ProSitePro files | PS50262 | G-protein coupled receptors family 1 profile. | 48 - 321 | - |
| Afu004794 |  |  |  |  |  |
| IPR000276 - G protein-coupled receptor, rhodopsin-like | PRINTS | PR00237 | Rhodopsin-like GPCR superfamily signature | 302 - 328<br>214 - 237<br>262 - 286<br>46 - 70<br>79 - 100<br>124 - 146<br>160 - 181 | $7.7 \times 10^{-51}$ |
|  | ProSitePro files | PS00237 | G-protein coupled receptors family 1 signature. | 130 - 146 | - |
| | Pfam | PF00001 | 7 transmembrane receptor (rhodopsin family) | 61 - 320 | $1.8 \times 10^{-68}$ |
| | SMART | SM01381 | 7TM_GPCR_Srsx_2 | 55 - 335 | $3.4 \times 10^{-8}$ |
| IPR000405 - Galanin receptor family | PRINTS | PR00663 | Galanin receptor signature | 152 - 168<br>110 - 128<br>93 - 107<br>289 - 309 | $1.2 \times 10^{-12}$ |
| IPR017452 - GPCR, rhodopsin-like, 7TM | ProSitePro files | PS50262 | G-protein coupled receptors family 1 profile. | 61 - 320 | - |
| Afu004211 |  |  |  |  |  |
| IPR000276 - G protein-coupled receptor, rhodopsin-like | Pfam | PF00001 | 7 transmembrane receptor (rhodopsin family) | 94 - 343 | $7.6 \times 10^{-17}$ |
| | PRINTS | PR00237 | Rhodopsin-like GPCR superfamily signature | 147 - 169<br>72 - 96<br>104 - 125<br>285 - 309<br>329 - 355 | $5.3 \times 10^{-05}$ |
| IPR017452 - GPCR, rhodopsin-like, 7TM | ProSitePro files | PS50262 | G-protein coupled receptors family 1 profile. | 87 - 347 | - |
| - | CDD | cd14978 | 7tmA_FMRFamide_R-like | 79 - 358 | $5.2 \times 10^{-34}$ |

Based on the search for conserved domains, we can conclude that all predicted sequences belong to 7 transmembrane and G-protein coupled receptors. All sequences except Afu017437 contain the Rhodopsin-like GPCR superfamily domain. However, the sequence of Afu004794 additionally contains a Galanin receptor domain. However, the sequence of Afu004794 additionally contains a Galanin receptor domain and Afu004211, Afu017437 an FMRFamide_R-like motif. Whereas exclusively opsin domains were found only in Afu005002 and Afu004575.

According to Gene Ontology terms determined using InterPro, all predicted *L. fulica* sequences belong to the G protein-coupled receptor signalling pathway (GO:0007186) except Afu005002, which additionally relates to phototransduction (GO:0007602) and visual perception (GO:0007601). By GO terms of PANTHER also all sequences belong to GO:0007186. Afu005002 additionally belongs to cellular response to light stimulus (GO:0071482) and phototransduction (GO:0007602), and Afu004575 belongs to cellular response to light stimulus (GO:0071482) and phototransduction (GO:0007602). Multiple sequence alignments were performed on the identified sequences containing conserved opsin domains (pfam00001 and cd00637), along with prediction of their 3D structures (Figure 1, Figure S1).

As a result, based on the analysis of multiple alignment results, it can be concluded that there is high variability and variability in the predicted sequences and functional opsin domains.

### BLAST analysis

BLAST analysis was performed to determine the percentage of identity of the predicted *L. fulica* sequences with other molluscan sequences (Table 3).

**Table 3.**
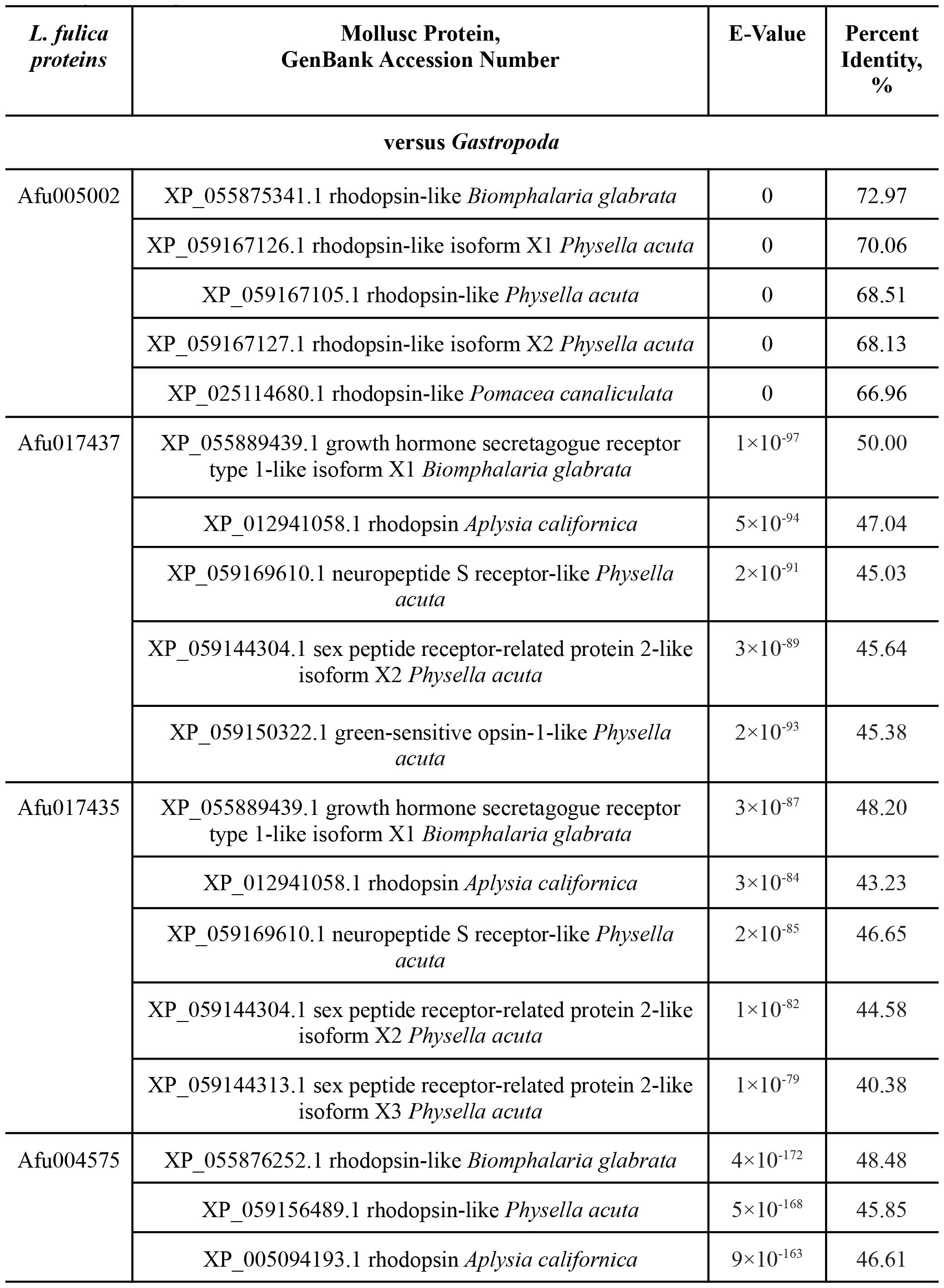
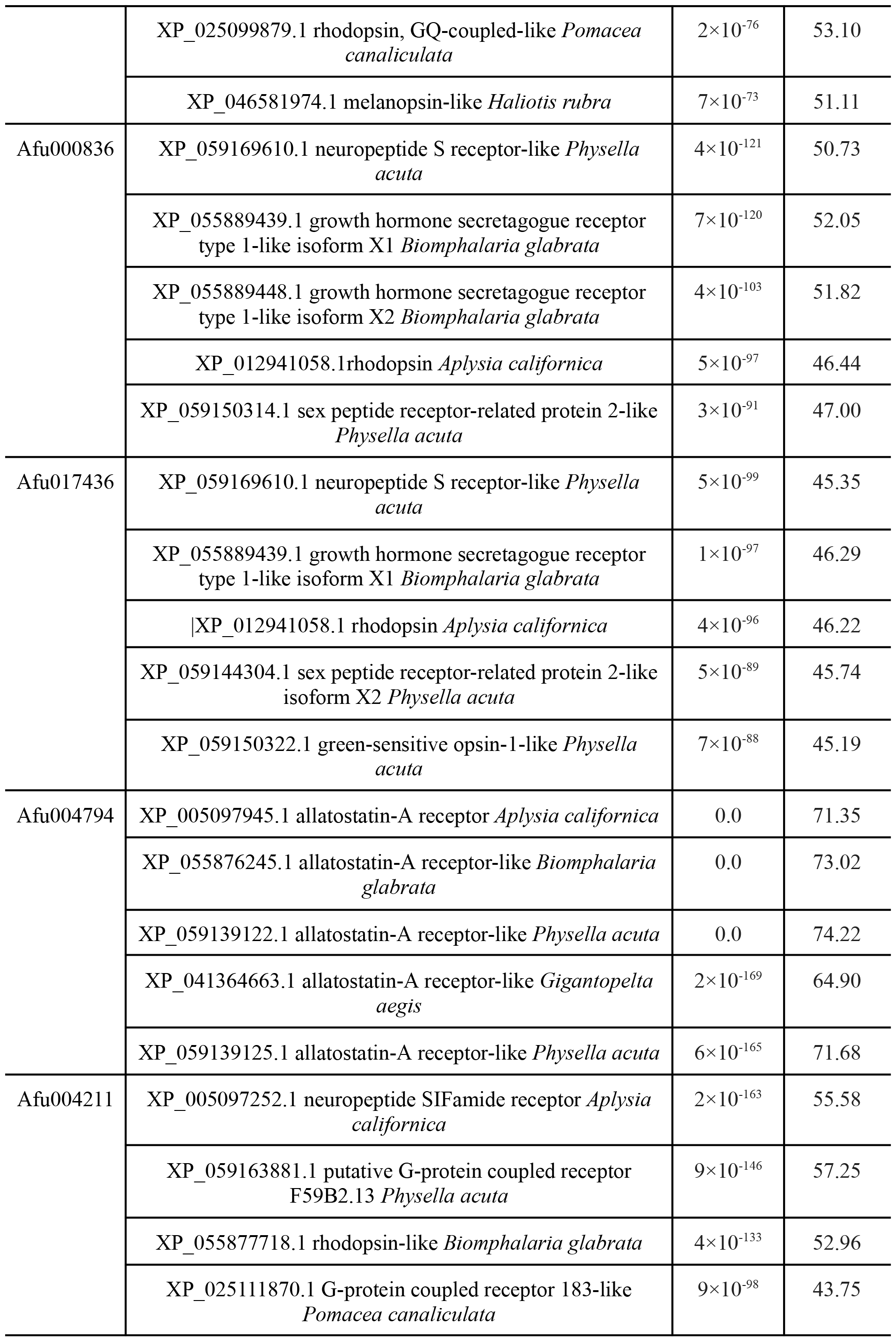
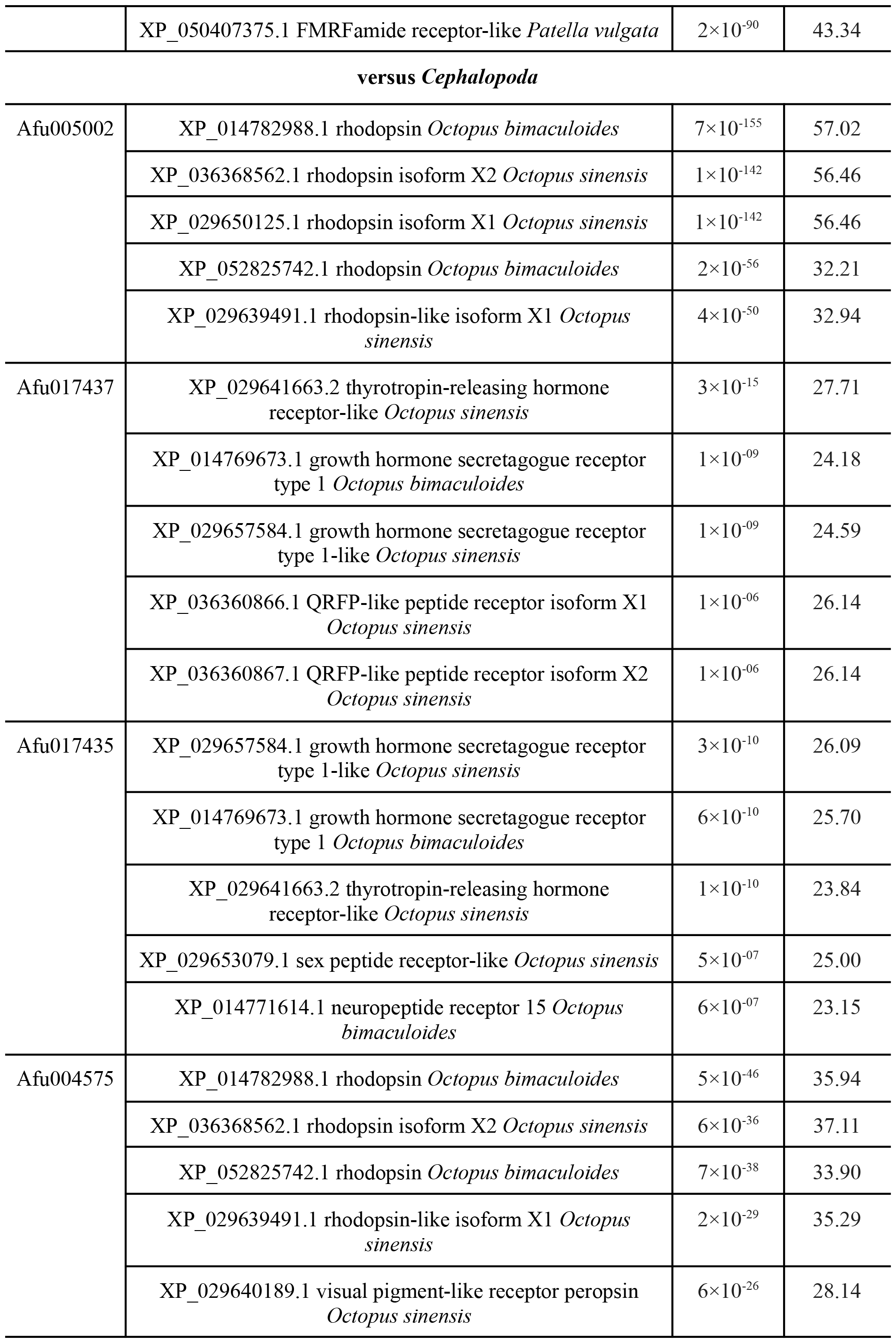
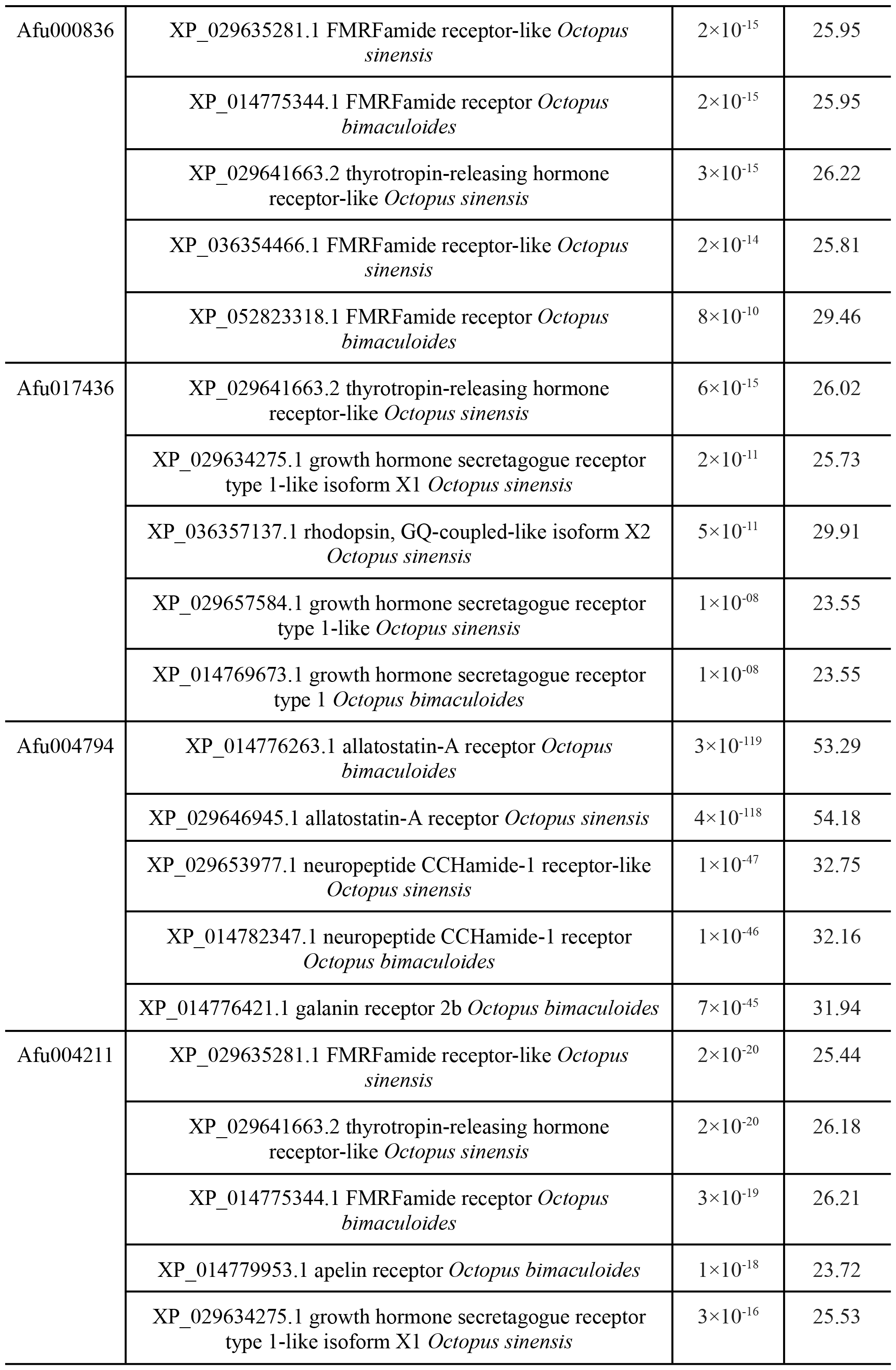
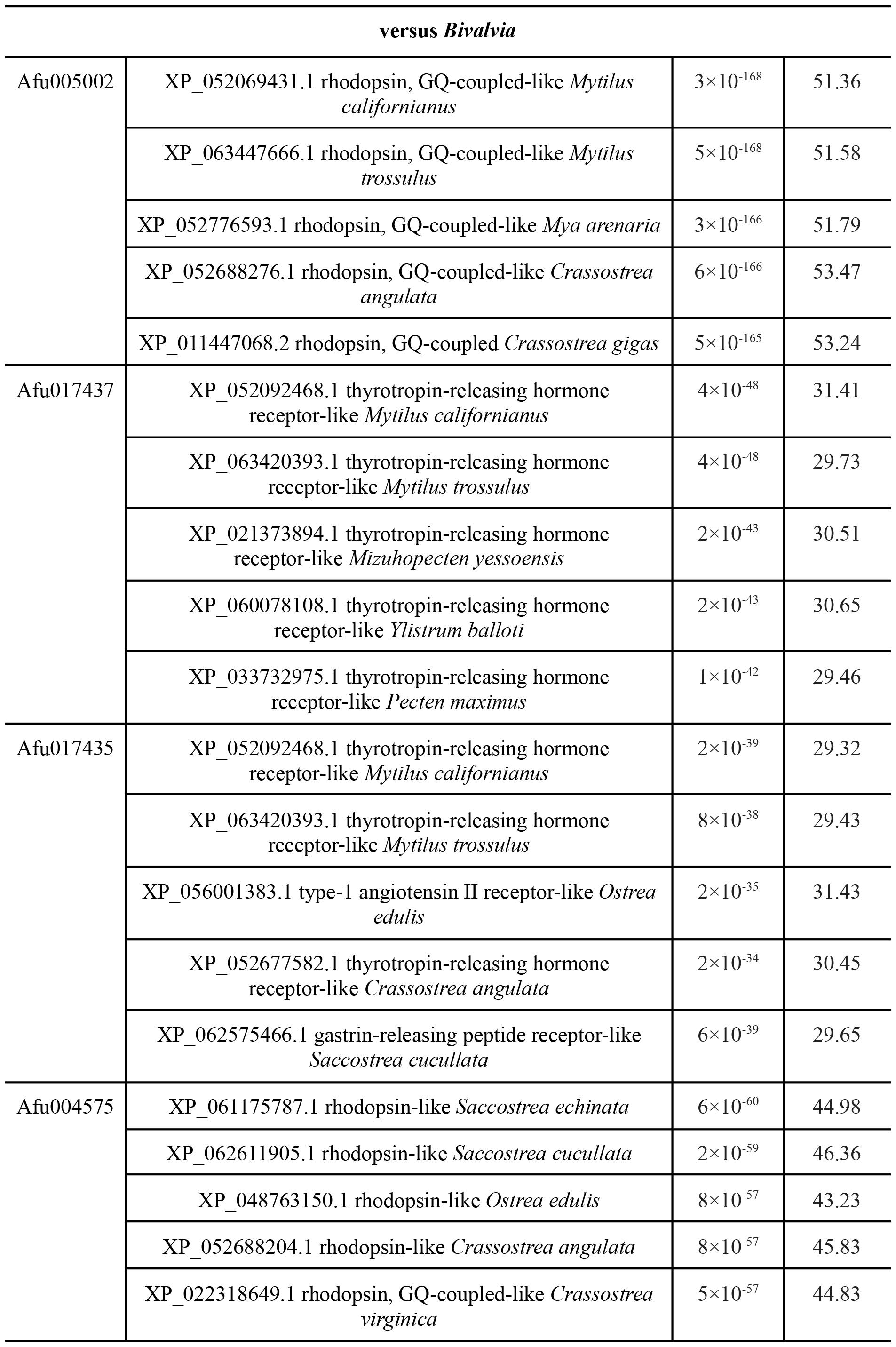
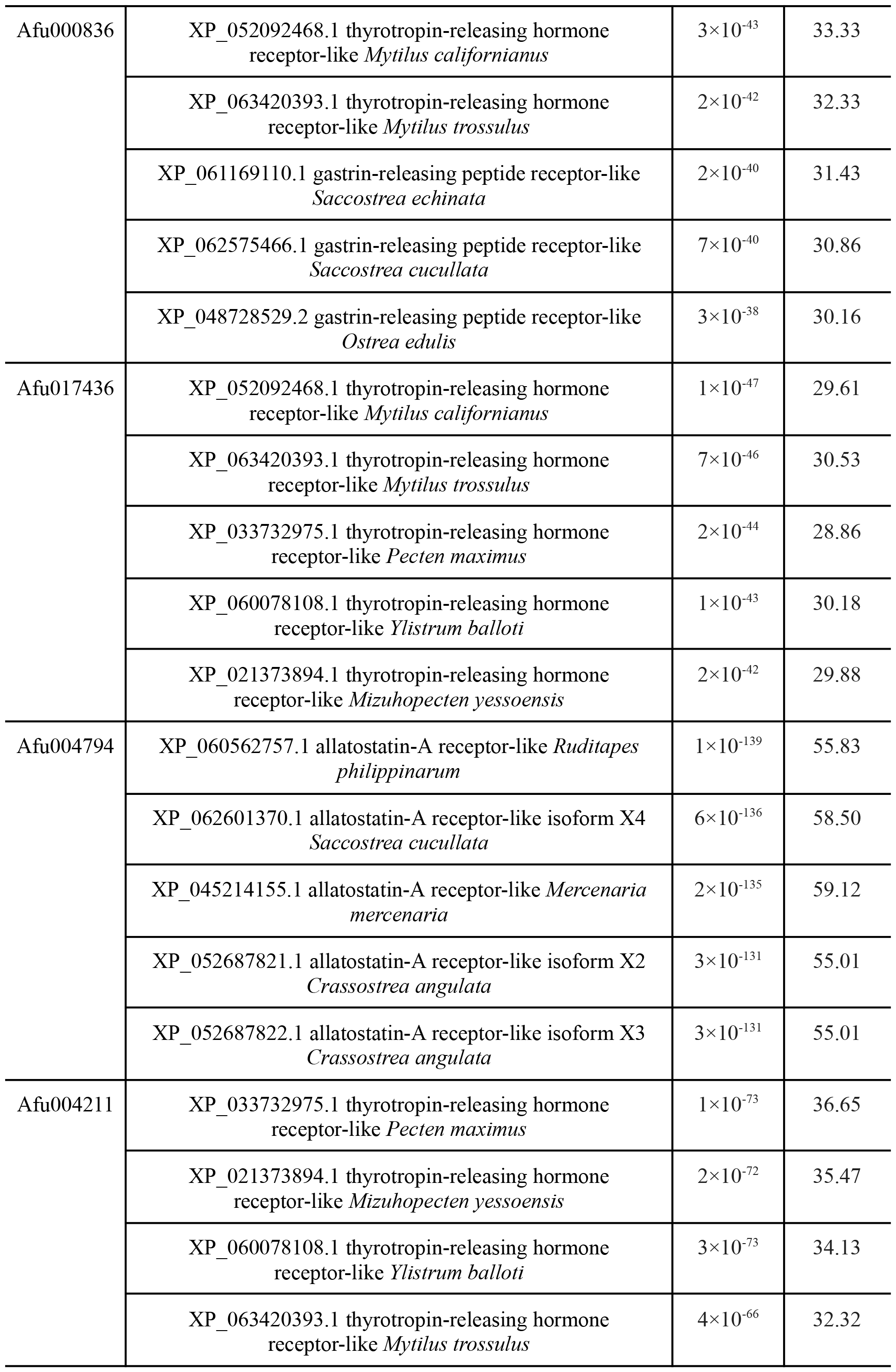
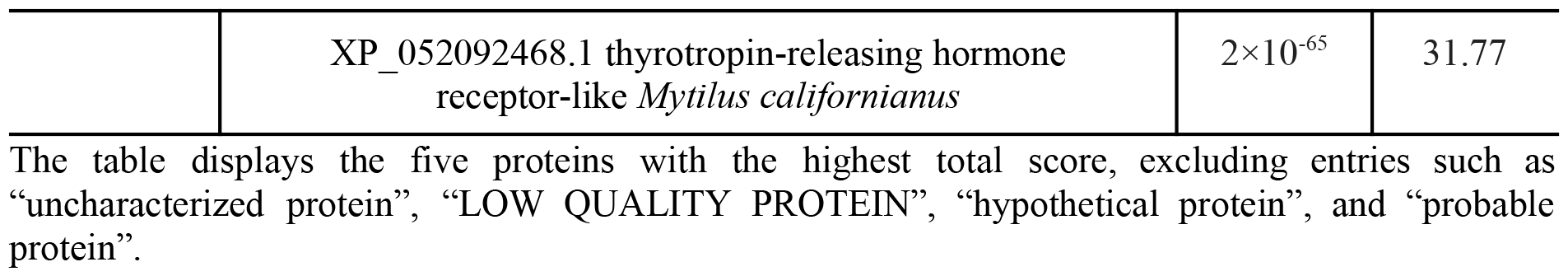
Percent identity of *L. fulica* predicted proteins versus *Gastropoda, Cephalopoda* and *Bivalvia* mollusc proteins by BLAST.

The table displays the five proteins with the highest total score, excluding entries such as “uncharacterized protein”, “LOW QUALITY PROTEIN”, “hypothetical protein”, and “probable protein”.

BLAST analysis as well as InterPro confirmed potential opsin affiliation for only two predicted proteins, Afu005002 and Afu004575, whereas all other sequences are only members of the 7-transmembrane receptor family.

### Structure alignment

Structural alignment was performed using PyMol for Afu005002 and Afu004575 sequences (Figures 3 - 4), which are the most likely candidates for the role of visual opsins in *L. fulica* based on the data obtained above (Figure 2 and Tables 1 - 2).

**Figure 2.**
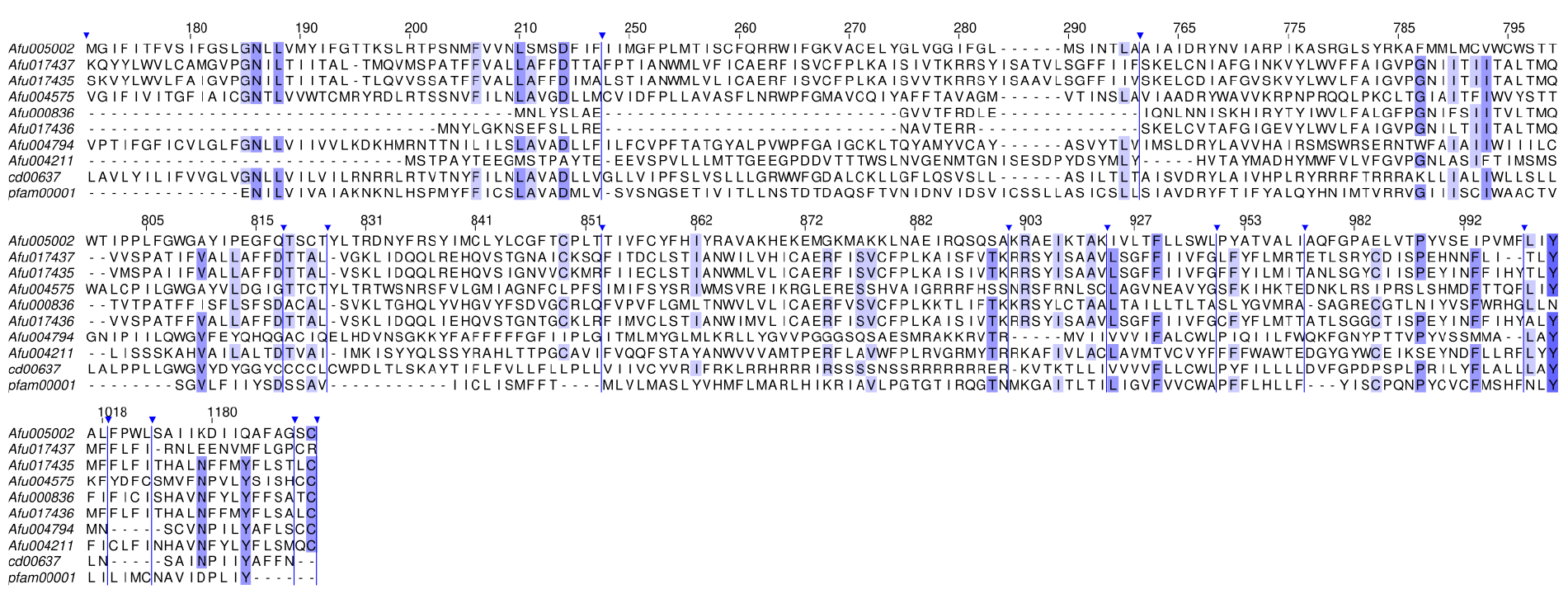
Alignment of amino acid sequences of *L. fulica* predicted proteins and opsin domains (pfam00001 and cd00637). The most conserved regions are presented. Multiple sequence alignments were performed by the Clustal Omega algorithm and refined with the MUSCLE algorithm. Amino acid residues with >60% identity are highlighted in lilac colour. The sign 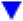 above the sequences indicates the excised regions. The single-letter amino acid designations correspond to the commonly used ones.

**Figure 3.**
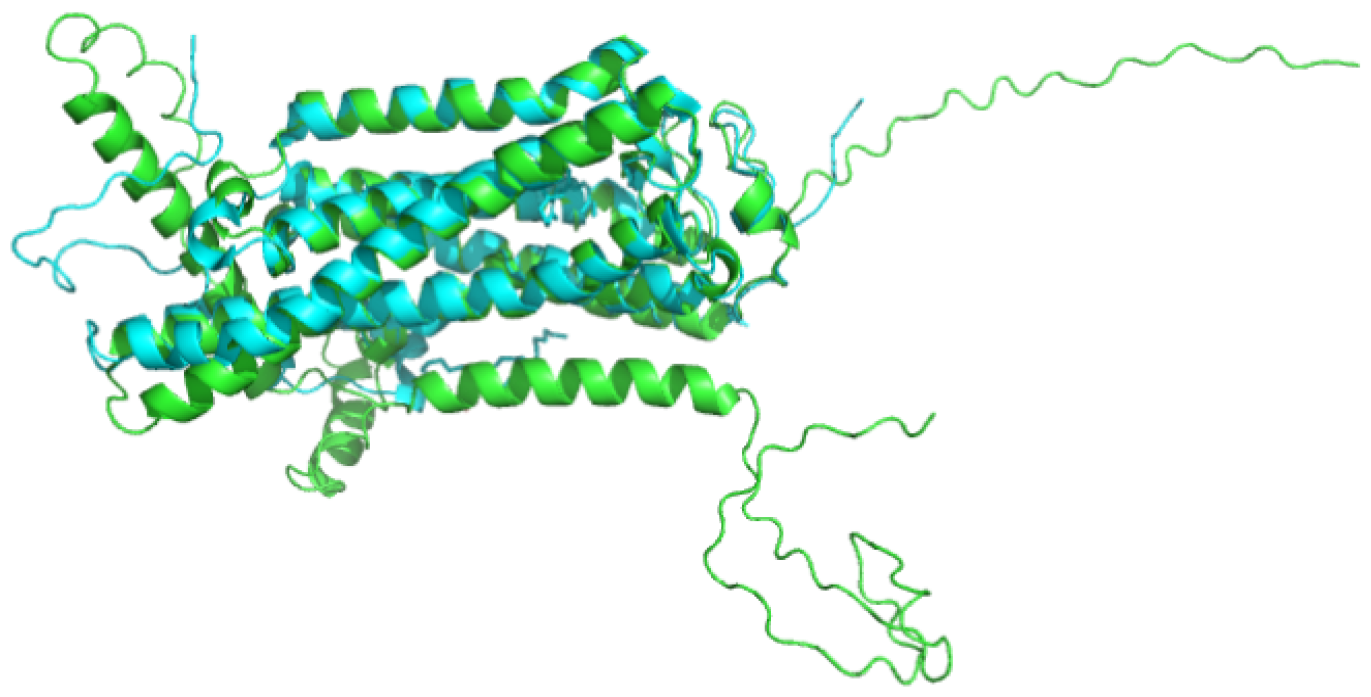
Alignment of 3D structures of the amino acid sequences of the Afu005002 *Lissachatina fulica* protein (green) with the rhodopsin of *Todarodes pacificus* (blue). The root mean square deviation (RMSD) of the two structures is 0.851 Å

**Figure 4.**
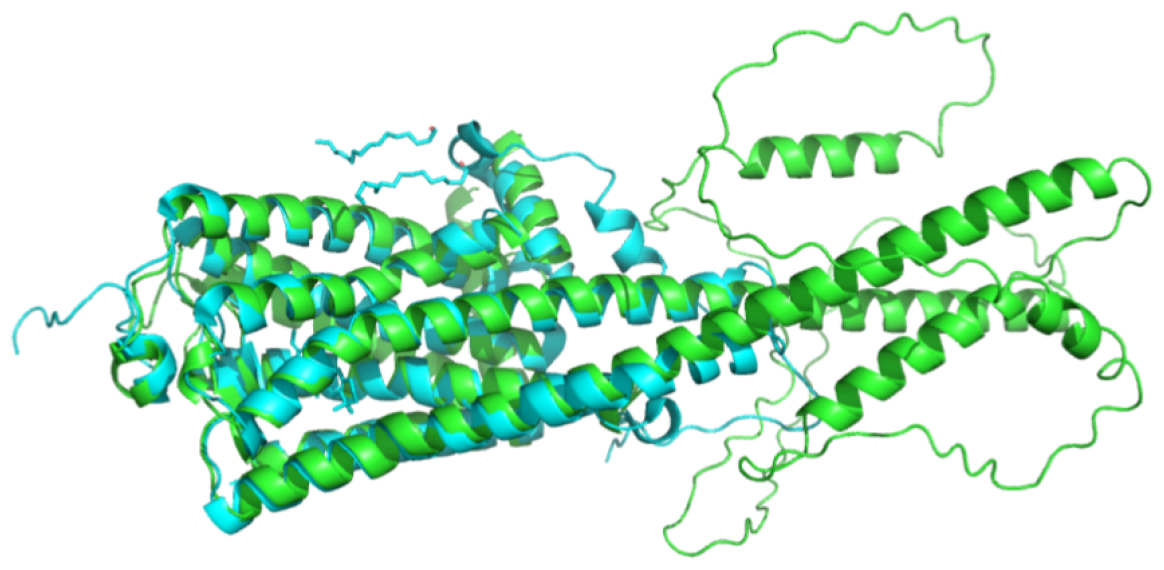
Alignment of 3D structures of the amino acid sequences of the Afu004575 *Lissachatina fulica* protein (green) with the rhodopsin of *Todarodes pacificus* (blue). The root mean square deviation (RMSD) of the two structures is 1.209 Å

Alignment of Afu004575 and Afu005002 with *L. fulica* protein indicates that the smallest value of the root mean square deviation (RMSD) of the predicted structures from the 3D structure of the squid *Todarodes pacificus* rhodopsin is 0.851 Å. This value belongs to the pairing of the squid rhodopsin with Afu005002. Based on the data presented above, we conclude that the most likely candidate for the role of visual opsin in *L. fulica* is the amino acid sequence Afu005002, which was taken for spectroscopic investigation analysis.

### Simulation of one-photon absorption spectrum

In this section we are going to present spectroscopic properties of 3D structure, predicted with AlphaFold2 based on the Afu005002 sequence. In order to estimate the effect of molecular environment under different pH level on absorption spectrum we consider several molecular models:

1. retinal (Fig.5A)
2. retinal and side chain of GLU175 counterion (Fig.5B)
3. retinal and protonated side chain of GLU175 counterion (Fig.6)

Since the initial orientation of the counterion proton can be different we have considered four different initial proton orientations, as it was done by Rozenberg and co-authors [37] and Emiliani and co-authors [38] (see Fig.5). Each orientation will be considered as an independent protonation pattern: pattern 3a, pattern 3b, pattern 3c, and pattern 3d (Fig.5,7).

#### Model 1 and Model 2

The structures of both models are presented in Figure 5. The optimised structure of Model 1 does not show significant changes with respect to the initial one. The computed excitation wavelength for Model 1 is red-shifted with respect to the recorded spectrum by Tamamaki [39.] for more than 50 nm. In contrast, the excitation wavelength computed for Model 2 shows a significant blue shift with respect to experimental data (∼150 nm). Moreover, optimization of retinal structure with deprotonated counterion shows a proton transfer from protonated Schiff base (pSB) towards the carboxylate oxygens of GLU175. Deprotonation of pSB leads to the strong blue shift of the main absorption band. For both models the first transition shows non-zero intensity, but the intensity of Model 2 is bigger by half with respect to Model 1 (Tab.4).

**Table 4.** The first excitation wavelengths for Model 1 and Model 2, as well as corresponding. The oscillator strength is given in parentheses. Experimental data was recorded at pH=7.4.

|  | Model 1 | Model 2 | Experiment |
| --- | --- | --- | --- |
| Wavelength, nm | 535.0 (0.95) | 329.0 (1.58) | ~480.0 |

**Figure 5.**
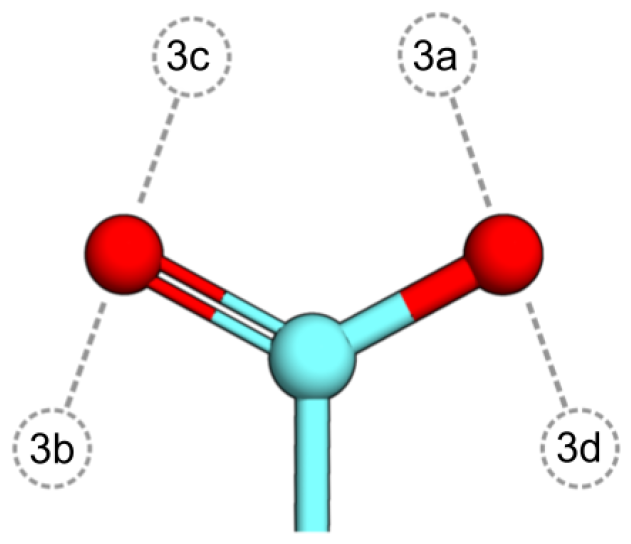
Four initial proton orientations of GLU175 counterion considered in this study. The proton is placed in the plane of the carboxyl group of GLU175.

**Figure 6.**
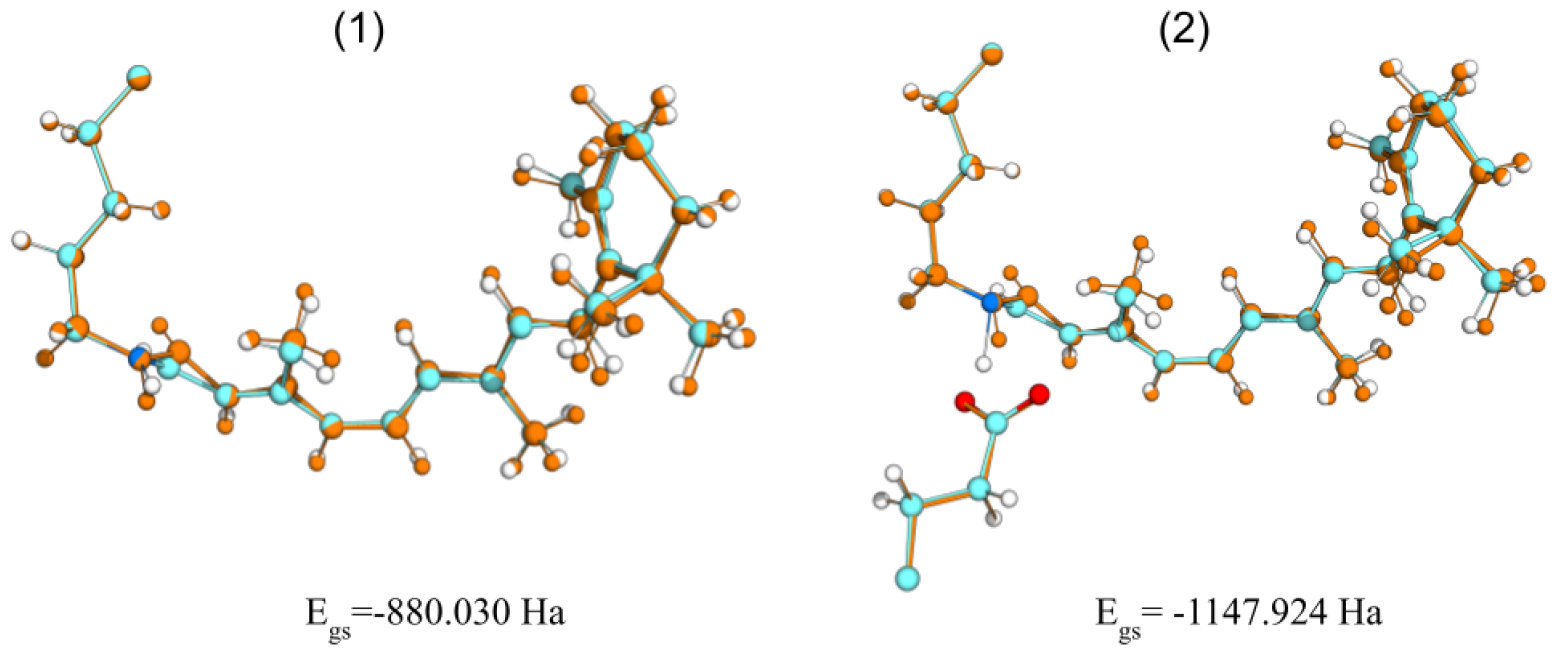
Structure of Model 1 (only retinal) and Model 2 (retinal + deprotonated counteroin GLU175) before (orange) and after (cyan) QM/MM optimisation. Ground state energy is presented at the bottom of the panels.

#### Model 3

Optimisation of protonation patterns does not show significant structural reorganisation except the rotation of the carboxyl group of GLU175 counterion. Generally, all protonation patterns of Model 3 show red-shifted excitation wavelength w.r.t. experimental spectrum. At the same time, patterns vary in both values of excitation wavelength and ground state energy. The best agreement with experimental spectrum was shown by 3d pattern, while patterns 3a and 3c indicate a large red shift with respect to recorded data: 31.4 nm and 48.3 nm, respectively (Tab.5). As a result protonation pattern 3b represents the most “successful” protonation model since it is the most stable structure and also shows a good agreement with the experimental data (Tab.5, Fig.7).

**Table 5.** The first excitation wavelengths computed for protonation patterns 3a, 3b, 3c, and 3d. The oscillator strength is given in parentheses. Experimental data was recorded at pH=7.4.

|  | 3a | 3b | 3c | 3d | Experiment |
| --- | --- | --- | --- | --- | --- |
| Wavelength, nm | 511.4 (1.1) | 502.7 (1.0) | 528.2 (1.0) | 501.1 (1.01) | ~480.0 |

**Figure 7.**
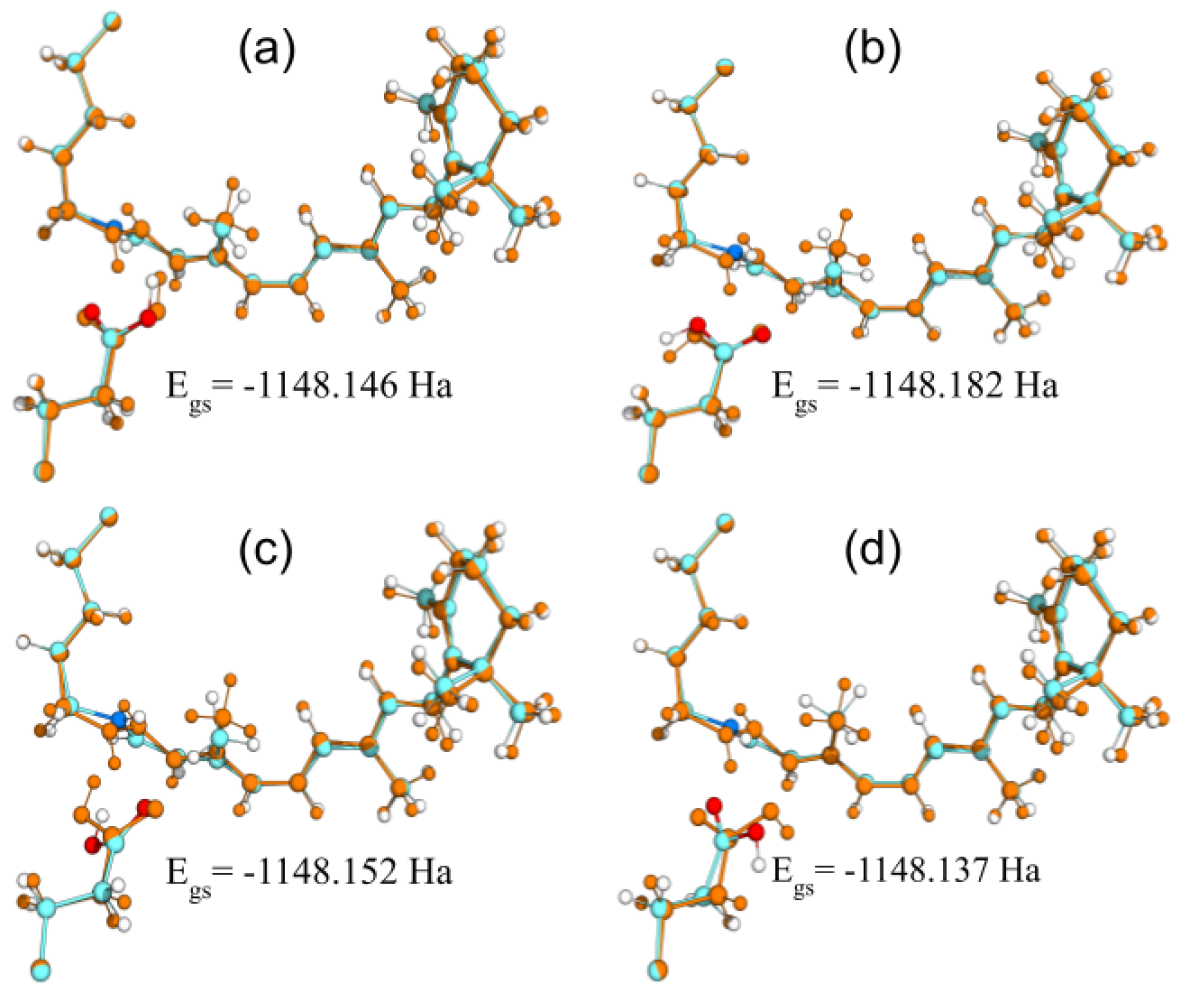
Structures of model 3 (retinal + protonated GLU175 counterion) with different initial position of the proton before (orange) and after (cyan) QM/MM optimization. Ground state energy is presented at the bottom of the panels.

## Conclusions

We have presented a complex theoretical study of visual opsin of *Lissachatina fulica*. Based on conformational search, bioinformatics and structural analysis the Afu005002 sequence is a promising candidate for a visual opsin of *Lissachatina fulica*. The computational study of spectroscopic properties 3D structure predicted based on Afu005002 also confirms the previous findings.

